# Modeling Epidemics in Seed Systems to Guide Management Strategies: The Case of Sweetpotato in Northern Uganda

**DOI:** 10.1101/107359

**Authors:** K. F. Andersen, C. E. Buddenhagen, P. Rachkara, R. Gibson, S. Kalule, D. Phillips, K. A. Garrett

**Affiliations:** Plant Pathology Department, University of Florida, Gainesville, FL 32611-0680, United States; Institute for Sustainable Food Systems, University of Florida, Gainesville, FL 32611-0680, United States; Emerging Pathogens Institute, University of Florida, Gainesville, FL 32611-0680, United States; Department of Rural Development and Agribusiness, Gulu University, Gulu, Uganda; Natural Resource Institute, University of Greenwich, United Kingdom

## Abstract

Seed systems are critical for deployment of improved varieties, but also serve as major conduits for the spread of seed-borne pathogens. We evaluated the structure of an informal sweetpotato seed system for its vulnerability to the spread of epidemics, and its utility for disseminating improved varieties. During the 2014 growing season, vine sellers were surveyed weekly in the Gulu Region of Northern Uganda. Our analysis draws on tools from network theory to evaluate the potential for epidemic spread in this region. Using empirical seed transaction data and estimated spatial spread, we constructed a network of seed and pathogen movement. We modeled the introduction of a pathogen, and evaluated the influence of both epidemic starting point and quarantine treatments on epidemic progress. Quarantine of 30 out of 99 villages reduced epidemic progress by up to 66%, when compared to the control (no quarantine), over 20 time steps. The starting position in the network was critical for epidemic progress and final epidemic outcomes, and influenced the percent control conferred by quarantine treatments. Considering equal likelihood of any node being an introduction point for a new epidemic, villages of particular utility for disease monitoring were identified. Sensitivity analysis identified important parameters and priorities for future data collection. The efficacy of node degree, closeness, and eigenvector centrality was similar for selecting quarantine locations, while betweenness had more limited utility. This analysis framework can be applied to provide recommendations for a wide variety of seed systems.

## Introduction

Seed systems, both formal and informal, are a critical component of global food security, but often also serve as human-mediated pathways for the regional and global dispersal of plant pathogens. Efforts to implement seed systems that work better for smallholder farmers in low-income countries have often been unsuccessful (Gibson and Kreuze 2015; Thomas-Sharma et al. 2016). Improving seed security – defined as timely access to quality planting material by all, at a fair price (Almekinders et al. 1994; Gibson et al. 2011; McGuire and Sperling 2013; Sperling 2008) – is vital for improved livelihoods, particularly for smallholder farmers. The provision of ‘clean’ or ‘pathogen-free’ seed is a major challenge to any seed system. In most low-income countries, seed systems are local and largely of unknown quality, with a majority of farmers keeping seed from previous seasons, or obtaining seed from neighbors, local traders, or local markets, with some instances of long distance trade (Gildemacher et al. 2009; Pusadee et al. 2009). Having robust analytic models of seed systems supports policy development and risk assessments against possible system disruption caused by system shocks such as novel pathogen or pest introduction, climate change, or political unrest.

Informal seed systems, without healthy seed certification protocols, may be at higher risk for epidemic introduction and spread. Newly introduced viruses can be particularly severe, as methods for detection may be limited or unavailable, and host resistance may be unattainable for several years. In this study, we consider an informal sweetpotato seed system, where ‘seed’ is not true botanical seed, but vine cuttings. In these types of vegetatively propagated seed systems, viruses and other seed-transmitted diseases introduce important risks to yield and quality degeneration over successive cycles of propagation, and methods of control are limited (Thomas-Sharma et al. 2016). It is important, therefore, that the risk of novel pathogen introduction and potential epidemic dynamics be understood for the swift recommendation of intervention (such as sampling, quarantine, variety deployment, and education).

The problem of seed-transmitted viral introduction was illustrated in 2014 when *Sri Lankan cassava mosaic virus* (causing cassava mosaic disease) was first detected in Southeast Asia, presumably being introduced through infected seed material. Efforts are still ongoing to mitigate spread and deploy resistance before this yield-robbing disease takes hold (Graziosi et al. 2016; Wang et al. 2016). Another dramatic example occurred in 2011 when maize lethal necrosis (MLN) was first reported in Kenya (Wangai et al. 2012) and soon was detected in several other Sub-Saharan African countries. MLN symptom expression results from coinfection with *Maize chlorotic mottle virus* (MCMV) and a potyvirus (Mahuku et al. 2015). Since its introduction, MLN has been detected in several East African countries including Ethiopia, Uganda, South Sudan, Tanzania, DRC, and Rwanda, and up to 22% yearly yield loss has been reported (De Groote et al. 2016; Hilker et al. 2017; Mahuku et al. 2015).

Network analysis is a powerful analytic tool, used across many disciplines, with recent advances in its utility for analyzing epidemic spread in human, animal, and plant systems (Shaw and Pautasso 2014; Silk et al. 2017). Seed systems are amenable to network analysis because they are inherently networks with a suite of actors (network nodes) that move both genetic material and information through space and time (dynamic or static network links) (Pautasso 2015). Until recent years, plant disease epidemiologists have given relatively little attention to the study of seed system (and plant trade) networks, although the movement of planting material plays a fundamental role in the spread of plant disease and the persistence of epidemics (Buddenhagen et al. 2017; Garrett et al. 2017; McQuaid et al. 2017; Nelson and Bone 2015; Pautasso 2015; Pautasso and Jeger 2008). Increasing availability of computational tools, coupled with advances in network analysis in the medical and social sciences, makes the implementation of network analysis for plant disease epidemiology more obtainable.

Network analysis can be used to evaluate nodes important for surveillance and mitigation of the movement of pathogens or other contaminants through seed systems and landscapes (Sutrave et al. 2012). To accomplish this, network statistics, such as measures of node betweeness, closeness, and degree centrality can be calculated to define key nodes and actors in a system, and forecast the risk of pathogen introduction, pathogen spread, or technology diffusion in a cropping system (Garrett 2012; Harwood et al. 2009; Moslonka-Lefebvre et al. 2011; Pautasso 2015; Pautasso and Jeger 2008; Sanatkar et al. 2015). For example, a study of wheat grain movement in the United States and Australia was implemented to identify key locations that could be strategically targeted for sampling and management of mycotoxins (Hernandez Nopsa et al. 2015). For the U.S. soybean rust epidemic, the utility of different selection methods was compared for targeting geographic nodes for sampling to forecast epidemic movement (Sanatkar et al. 2015). Although there are a number of statistics available that are likely to be associated with the importance of a node in an epidemic, it is an open question as to which statistics are most important for prioritizing nodes for monitoring and management in real-world networks (Holme 2017).

Previous studies of seed systems have focused on understanding the effects of social ties and how well seed system networks may conserve variety diversity in the landscape (Pautasso 2015; Pautasso et al. 2013). Abay et al. (2011) applied network analysis in a study of informal barley seed flows in the Tigray region of Ethiopia, with the goal of informing breeding or technology deployment strategies. Network properties, such as betweenness and degree centrality, were used to characterize key nodes and their role in connecting the seed system.

These metrics measure the importance of a node in terms of the number of connections it has, and the number of paths across the network of which it is a part, respectively. Pautasso et al. (2015) further analyzed this empirical seed network data, finding that the degree distribution of the network, and particularly the out-degree of the starting node, influenced the percentage of the network that could be reached by a new variety.

Our study draws on concepts developed for studying epidemic spread in hypothetical and empirical large- and small-scale plant trade networks (Buddenhagen et al. 2017; Moslonka-Lefebvre et al. 2011; Nelson and Bone 2015; Pautasso 2015; Pautasso and Jeger 2008; Pautasso et al. 2010). We consider theory about the influence of node in- and out-degree on variety dissemination (and pathogen transmission) in small-scale, real-world seed networks (Pautasso 2015). We also expand on concepts developed to assess quarantine influence on disease spread in hypothetical trade networks (Nelson and Bone 2015) in a new set of simulation experiments, where quarantine nodes were not selected after initial pathogen detection, but based on their network centrality measures. Here we define quarantine as the restriction of the exchange of infected seed material, likely though phytosanitary regulation. We further explore the question of quarantine efficacy based on the epidemic starting point and the chosen centrality predictor.

We model epidemic spread and mitigation in a landscape of farming villages, where a portion of the seed transaction structure is based on geo-spatial proximity of villages. Our model of the contact structure (links) between villages is based on the much higher tendency for farmers to exchange planting material with neighboring villages than with those that are distant (Perales et al. 2005; Pusadee et al. 2009). We also consider the sensitivity of parameter choices in our model and their impacts on epidemic outcomes. Improvement of these analytic tools is particularly important for understanding the epidemiology of pathogen spread in vegetatively propagated crops, where social contact structures via the exchange of planting material is a major driver of epidemics.

Seed systems are a suite of actors, or nodes, including farmers, multipliers, traders, NGOs, seed companies, breeding organizations, and communities. The connections between these nodes representing formal and informal interactions, are complex and require analyses that address this complexity. Network analysis allows us to simulate multiple scenarios in known systems, including the impact of epidemic spread or management deployment. In this study we propose a general framework for analyzing such networks that can be translated to a broad range of seed systems. In this paper, we, i) analyze, as a case study, key network properties within a seed system important to regional food security; ii) evaluate variety dissemination within the network, comparing the distribution of higher-nutrient introduced varieties and landraces; iii) model the progress of a potential seed-borne pathogen introduced into the network, and compare the use of different network statistics as selection criteria for quarantine nodes; and iv) perform sensitivity / uncertainty analysis to determine the influence on epidemic outcomes of the parameters used to construct the village-to-village transaction networks.

## Materials and Methods

### Study System: Sweetpotato in Northern Uganda

This study examines sweetpotato vine transactions in Northern Uganda. Sweetpotato is a major staple food crop in many African countries, and Uganda is the second largest producer in Africa, fourth globally (FAOSTAT 2013). Sweetpotato is generally grown by women in Uganda in small plots of land, close to the household, and is important for household food security (Behrman 2011; Johnson and Gurr 2016). In the last decade, sweetpotato has increased in importance due to the introduction of a β-carotene biofortified crop, Orange-fleshed Sweetpotato (OFSP), by HarvestPlus, part of the CGIAR Research Program on Agriculture for Nutrition and Health. OFSP varieties were introduced with the goal of addressing vitamin A deficiency in women and children in this region (Behrman 2011).

Viral diseases are major biotic limiters to sweetpotato production in Uganda and throughout Sub-Saharan Africa, with the most yield-limiting being sweet potato virus disease (SPVD), which occurs when a plant is co-infected with *Sweet potato feathery mottle virus* (SPFMV) and *Sweet potato chlorotic stunt virus* (SPCSV) (Karyeija et al. 1998). Seed degeneration is the successive loss in yield over generations of vegetatively-propagated seed planting material due to the accumulation of viruses and other seed borne pathogens (Thomas-Sharma et al. 2016). Degeneration is highly problematic in informal seed systems where farmers tend to save seed season-to-season and where certified seed sources are rare or non-existent. The normal planting material for sweetpotato is foliar cuttings, and both SPFMV and SPCSV can be transmitted to succeeding generations this way, with evidence of yield degeneration over five generations in high pressure fields in Uganda (Adikini et al. 2015). SPVD has not yet been reported in Northern Uganda, likely because the extended dry season in this region is often unfavorable for the whitefly vector (Richard Gibson, *personal observation*). Changes in climate patterns or vector range, however, could potentially extend the range of this disease into this region. The prospect of novel pathogen introduction makes it important to model epidemic scenarios to inform intervention strategies.

In Northern Uganda, sweetpotato seed material is sold in small bundles of vine cuttings. Distribution of these vine cuttings is largely informal, consisting of smallholder farmers who have access to fields with adequate moisture to produce roots and vines through the extended dry season, which typically lasts from December to April (Gibson 2013). These off-season multipliers generally produce local landraces, which tend to be well-adapted white-fleshed cultivars. (Gibson 2013). Vine cuttings are not easily stored, and because of the single, extended dry season in Northern Uganda, vines need to be obtained by farmers at the beginning of each season (Gibson et al. 2011). There are also several formal institutions involved in sweetpotato breeding and distribution in Uganda, including the National Sweetpotato Program (NSP), the International Potato Center (CIP), private sector enterprises, and NGOs (Gibson 2013).

### Survey Methods

A survey of vine multipliers and sellers was conducted in 2013-2014 in the Gulu Region of Northern Uganda, fully described by Rachkara et al. (2017). A more complete cohort of multipliers and sellers was surveyed in 2014, the focus of our analysis. Each seller was visited weekly from the start of the growing season (April) and surveyed twice per week to record all transactions (purchases and sales) that occurred in the period since they were last visited until the end of the season (August). Volume of transaction (number of bundles), price, variety, origin of buyer, and buyer type (farmer or seller) were recorded. In this study, a small bundle refers to 50 vines cut to 40 cm in length. Large bundles are equal to 20 small bundles. Because of the high volume of transactions, names of individual buyers were not collected and therefore sales transactions were summarized by the buyer’s village.

### Seed Network Analysis

Nodes in this analysis include sellers and villages, with one set of directed links representing vine sales from an individual seller to an individual village (a bipartite graph). Villages in this region of Uganda typically are composed of 40-60 households. Although several transactions were recorded on a weekly basis, transactions were aggregated for this analysis so that seller-to-village links represent the existence of at least one transaction over the course of the season. Seller-village links are based entirely on the data from Rachkara et al. (2017).

### Incorporating village-to-village spread

To incorporate epidemic risk in the seed system outside of spread through sellers, we evaluated a second set of geo-spatially derived links representing village-to-village spread through informal exchange of planting materials or vector movement, composing a second network. The existence of village-village links was modeled using a power law function, and combined with transaction links to form an expanded adjacency matrix (Fig. 1). The power law function captures the tendency for seed exchange to be more likely between villages that are geospatially proximate. That is, farmers have a higher chance of exchanging seed with farmers in neighboring villages than with farmers in villages that are far away. A link represents a seed movement event (transaction) or movement of vectors that may be viruliferous. The power law equation used here was y = ad^−β^, where *d* is the Euclidean distance between a pair of villages, *y* is proportional to the probability of movement between the villages, and *a* is a scaling factor. As default values, we used a = 1 and β = 1.5. The seed transaction threshold (*t*) was set to 0.01, such that when y > 0.01 for two villages, a link was created between them. Village-to-village links were established once and the same underlying adjacency matrix was used across simulation experiments for a given parameter combination. In individual realizations, the underlying adjacency matrix was modified as links were maintained with a fixed probability, as described below. A sensitivity / uncertainty analysis was performed to determine the influence of the parameters β and *t* on the analysis. It is important to note that only villages of farmers who purchased vines from sellers surveyed in this study were included in this analysis, and there may be villages in this region that contribute to risk, but were not included.

**Figure 1.**
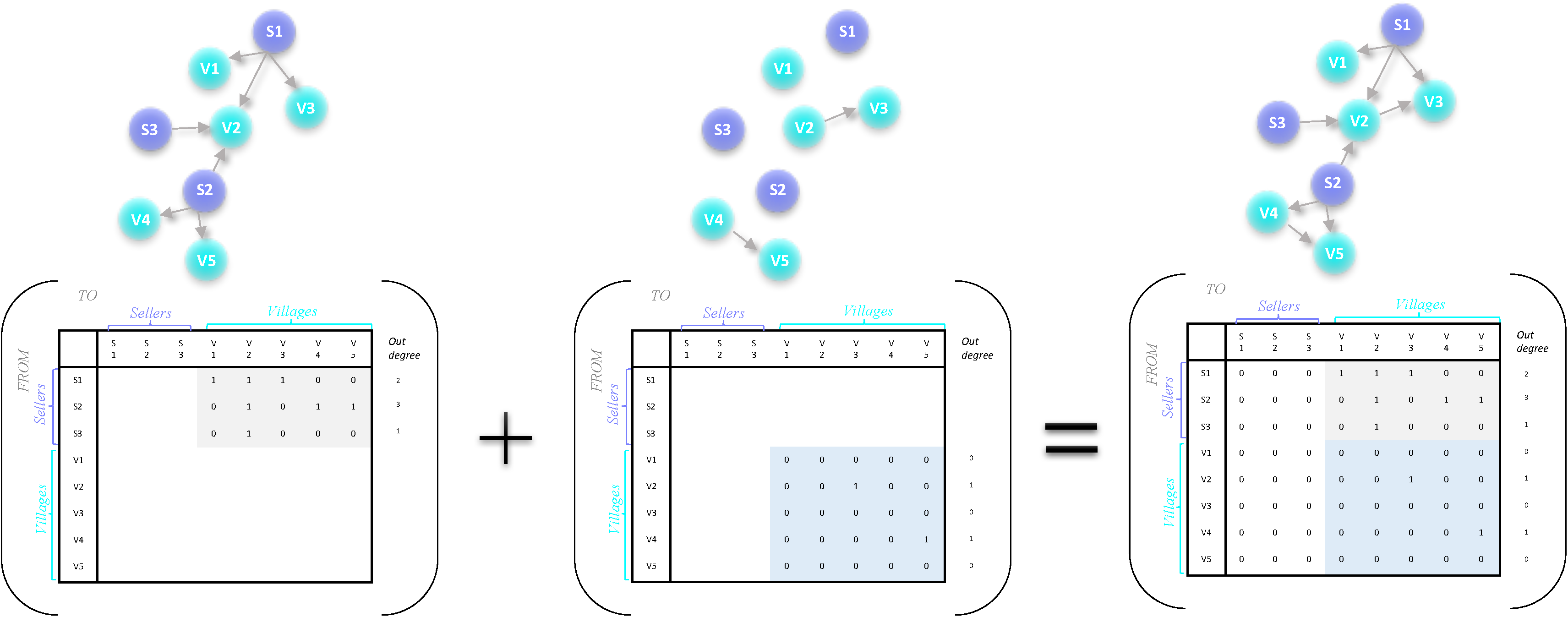
A schematic illustrating how two types of data were combined: A) seller-to-village links reported in the 2014 vine seller survey, and B) village-to-village links estimated based on the distance between villages using a power law function to estimate the probability of movement. The schematic represents a hypothetical network of three sellers and five villages, each represented as a node in the network. A link between nodes is represented as a 1 in the matrix, and absence of a link is indicated by a 0. Matrices were combined in C) a “complete network” with both seller-to-village and village-to-village links. Note that in this study all seller-to-seller links and village-to-seller links were set to zero, with no transactions taking place in this direction. A “complete network” based on the Ugandan sweetpotato data was used in the simulation experiments presented in this study.

Network properties, such as number of nodes and network density, were calculated for the 2014 season. Node measures such as degree, closeness, eigenvector, and betweenness centrality were evaluated for both villages and sellers (Table 1). All analyses were conducted in the R programing environment (R Core Team 2017) using custom code along with software packages including igraph (Csárdi and Nepusz 2006).

**Table 1.**
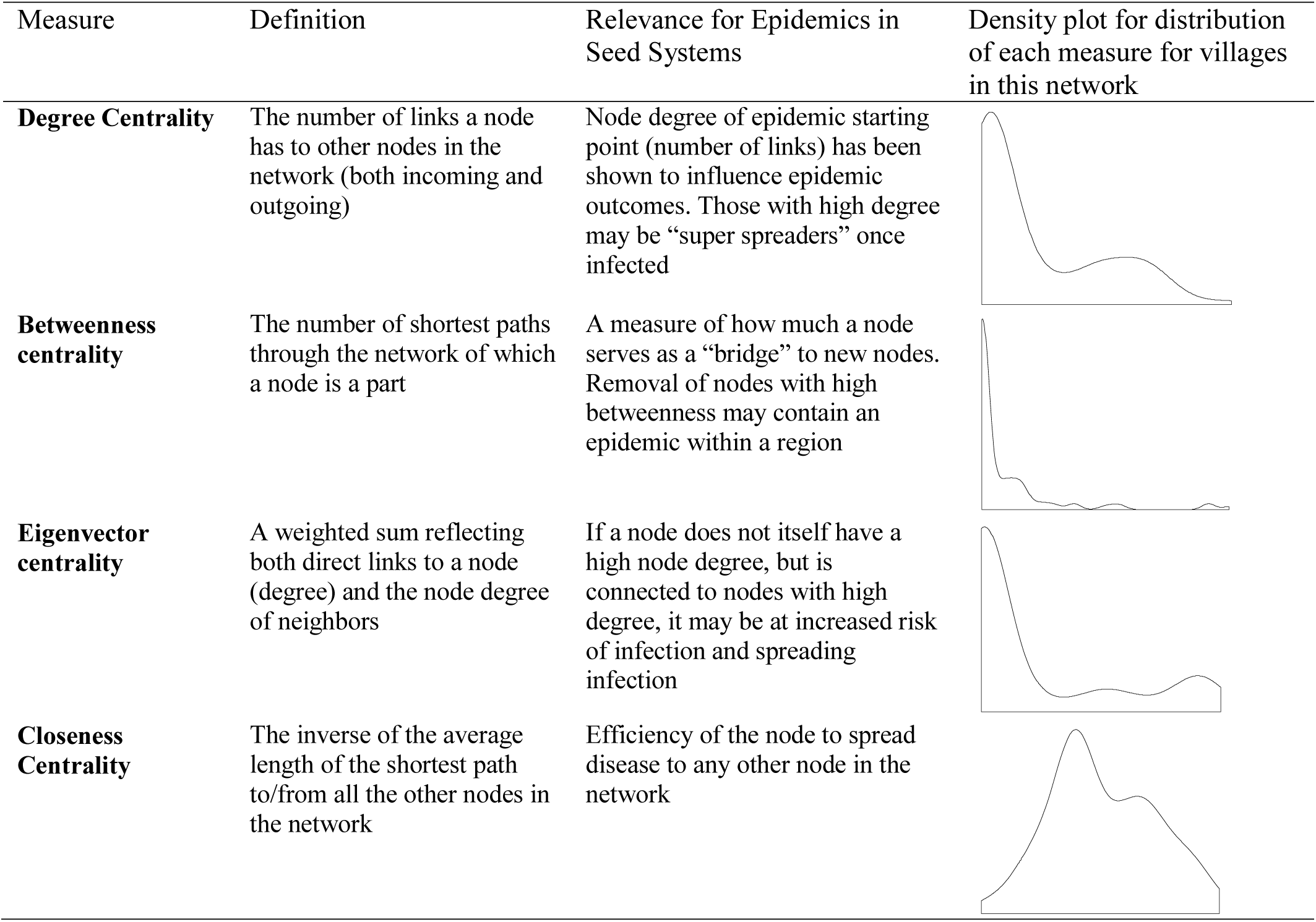
Common network node centrality measures

| Measure | Definition | Relevance for Epidemics in Seed Systems | Density plot for distribution of each measure for villages in this network |
| --- | --- | --- | --- |
| **Degree Centrality** | The number of links a node has to other nodes in the network (both incoming and outgoing) | Node degree of epidemic starting point (number of links) has been shown to influence epidemic outcomes. Those with high degree may be “super spreaders” once infected |  |
| **Betweenness centrality** | The number of shortest paths through the network of which a node is a part | A measure of how much a node serves as a “bridge” to new nodes. Removal of nodes with high betweenness may contain an epidemic within a region |  |
| **Eigenvector centrality** | A weighted sum reflecting both direct links to a node (degree) and the node degree of neighbors | If a node does not itself have a high node degree, but is connected to nodes with high degree, it may be at increased risk of infection and spreading infection |  |
| **Closeness Centrality** | The inverse of the average length of the shortest path to/from all the other nodes in the network | Efficiency of the node to spread disease to any other node in the network |  |

**Table 2.** Parameters describing dispersal probabilities in a seed system network

| Parameter | Definition | Values | Comments |
| --- | --- | --- | --- |
| $\beta$ | Exponent of the power law, where $\beta > 0$. Influences the shape of decay | 0.5-2 (0.5, 1.0, 1.5*, 2.0, 2.5) | Larger values of $\beta$ result in steeper decay, and fewer village to village links |
| $t$ | Seed transaction threshold. Applied to power law transformed distance data. | 0.001-0.15 (0.001, 0.005, 0.01*, 0.015, 0.02) | Influences how close neighboring villages need to be to have a link formed in the village-to-village network. |
| $P_t$ | Probability of transmission of infection at each timestep, over an established link | 0.05-0.15 (0.05, 0.01*, 0.015) | Influences the probability that infection will be transmitted along a network link in each time step, from seller to village, or village to village |

*\*indicates the value that was used as the default in simulation experiments*

### Modeling Epidemic Spread

This study draws on tools from graph theory to better understand the potential for epidemic spread within a real-world seed network. This analysis used both observed 2014 seed transaction data and estimated spatial transactions to simulate the invasion of a potential seed-borne pathogen. Utilizing the above described network, we performed simulation experiments to address the following questions. (1) What are the optimum risk-based surveillance locations for pathogen monitoring, if there is equal likelihood of pathogen introduction at each node? (2) What effect does seller at which disease is introduced have on disease progress and final disease outcome? (3) How much can disease spread be limited by implementing quarantine treatments, where quarantined nodes cannot become infected or spread disease? (4) How do network statistics compare for their utility for selecting quarantine locations? In each experiment, simulations were conducted over 20 timesteps and repeated 500 times. The probability of pathogen transmission (*P*_*t*_) occurring in each of the generated links was set to 0.10. The probability of persistence (*P*_*p*_; entries on the diagonal of the adjacency matrix) was set to 1.

### Experiment 1: The value of villages as risk-based surveillance locations

We evaluated the importance of each village as a risk-based pathogen surveillance location. In this scenario analysis, all nodes (sellers and villages) were assigned an equal likelihood of being the starting point for epidemic introduction. For each possible combination of an epidemic starting node and a sampling node, we determined the number of nodes that could become infected by the time the pathogen is detected at the sampling node, as in Buddenhagen et al. (2017). In this analysis we compare only villages, not sellers, for their relative value as locations for epidemic monitoring. In 500 realizations, the mean and variance of the number of nodes infected by the time the pathogen was detected in each village were evaluated, and each village was assigned a “surveillance score”. Summarizing over all the potential starting nodes allows for the comparison of the importance of each village as a potential location for monitoring, with each potential starting location equally likely. Simulations were implemented using custom R code (R core team 2017).

### Experiment 2: Modeling epidemic progress with variable starting sellers

The second simulation experiment evaluated the potential spread through the seed network, where the epidemic starts with a single seller that either sells only landraces (86% of sellers, Fig. 3) or sells only orange fleshed sweet potato varieties (14% of sellers, Fig. 3). The two sellers with the highest out-degree in each category (OFSP and landrace seller) were chosen for comparison in this analysis, representing the greatest risk for each type. The out-degree of the starting node is generally strongly correlated with final epidemic outcome (Pautasso 2015; Pautasso et al. 2010). At the start of each realization (Time 1), the “starting seller” had an infection status of 1 and all others were set at 0. The starting seller maintained this infection status throughout the realizations, and all other sellers remained uninfected. In each timestep, 10% of the links from the infected seller node resulted in an “infection event” where the status of the linked village node became 1. Villages that became infected after one time step maintained infected status in the subsequent time step (Time t+1) and could thus infect villages to which they had links with the same probability (*P*_*t*_ = 0.01) in the subsequent time step (Time t+2; probability of infection persistence, *P*_*p*_ = 1; see Supplemental Figure 1). Infection was evaluated across 20 timesteps in 500 realizations, to evaluate the frequency distribution of outcomes. Simulations were carried out using custom R code (R core team 2017). To evaluate the differences in disease progress between starting nodes, the area under the disease progress curve (AUDPC) was calculated by summing the trapezoids between timesteps under the curve.

### Experiment 3: The influence of quarantine on infection dynamics

We evaluated the influence of quarantine on disease spread over time. Quarantine was defined here as the removal of the potential for a node to transmit or acquire infection in each time step. The influence of 10, 15, 20, 25 and 30 quarantined nodes on infection spread over time was compared with a control in which there was no quarantine treatment. Quarantined nodes (villages only) were selected initially based on node degree, with villages with the highest node degree selected first. For consistency, the same two sellers (S_25 and S_15) that were chosen to start the epidemic in the previous experiment were used in this analysis. Simulations were implemented as described above, over 20 timesteps in 500 realizations, with the AUDPC calculated for each quarantine treatment. To further explore the influence of starting seller on quarantine treatment, we calculated quarantine efficacy, or the proportional reduction in disease as compared to the non-quarantined treatment, as (AUDPC_no quarantine_ –AUDPC_quarantine_) / (AUDPC_no quarantine_) for all 27 sellers.

We also compared the utility of key network node statistics (Table 1) for the selection of nodes as quarantine candidates. We performed the above quarantine analysis (for quarantine of 30 nodes, two starting sellers) with villages selected for quarantine based on their node betweenness centrality, closeness centrality, and eigenvector centrality, in comparison to a scenario where 30 villages were drawn at random (Table 1). The value of each villages as a risk-based surveillance location, calculated in Experiment 1, was also compared for its utility to select quarantine candidates. In the evaluation of each method for ranking the likely value of nodes for quarantine, nodes were ranked in importance based on that method and the top ranked 15, 20, 25 or 30 nodes were selected for quarantine.

### Sensitivity analysis and uncertainty quantification

To evaluate the influence of parameter choices for parameters describing system-specific and often unknown features such as the dispersal kernel, we evaluated the effects of varying parameter values on the epidemic outcome (summation of infection status of all nodes at the end of 20 timesteps) for each of the parameter combinations described in Table 1. Parameters *β* and *t* influence the number of links formed between villages, ultimately influencing the likelihood of transmission of a pathogen at each time step. The probability of transmission (*P*_*t*_) influences the percentage of links that will transmit infection in each timestep, ultimately influencing epidemic progress. These analyses were done for both infection starting at S_15 (the highest degree landrace seller) and S_25 (the highest degree OFSP seller).

## Results

### Network Properties

In 2014, 27 sellers were tracked, resulting in a total of 878 individual vine sales to buyers from 99 distinct villages. The seller-to-village portion of the adjacency matrix was estimated based on aggregated transactions from sellers to villages (Figure 2a). This network has a total of 126 nodes and 205 links (link density = 0.013). After the addition of village-to-village links (link density = 0.047, Figure 2b), the number of links increased to 743, roughly representing the number of potential vine transactions in each time step in simulation experiments. Node degree, or the number of incoming (in-degree) and outgoing (out-degree) links was calculated for both seller nodes (mean = 7.6, min =1, max =42) and village nodes (mean = 12.9, min = 1, max = 60; Supplemental Fig. 2). The degree distribution of this network indicates scale-free properties (Barabasi and Albert 1999), with a high number of nodes having few link and fewer nodes having many links. The 30 villages with the highest node degree (both in- and out-degree) were selected as candidates for quarantine in subsequent analyses. Betweenness, closeness, and eigenvector centrality were also measured for all villages in this analysis (Table 1) and used in subsequent analysis. These four measures of centrality were positively correlated based on Pearson’s correlation coefficient (Supplemental Fig. 3).

**Figure 2.**
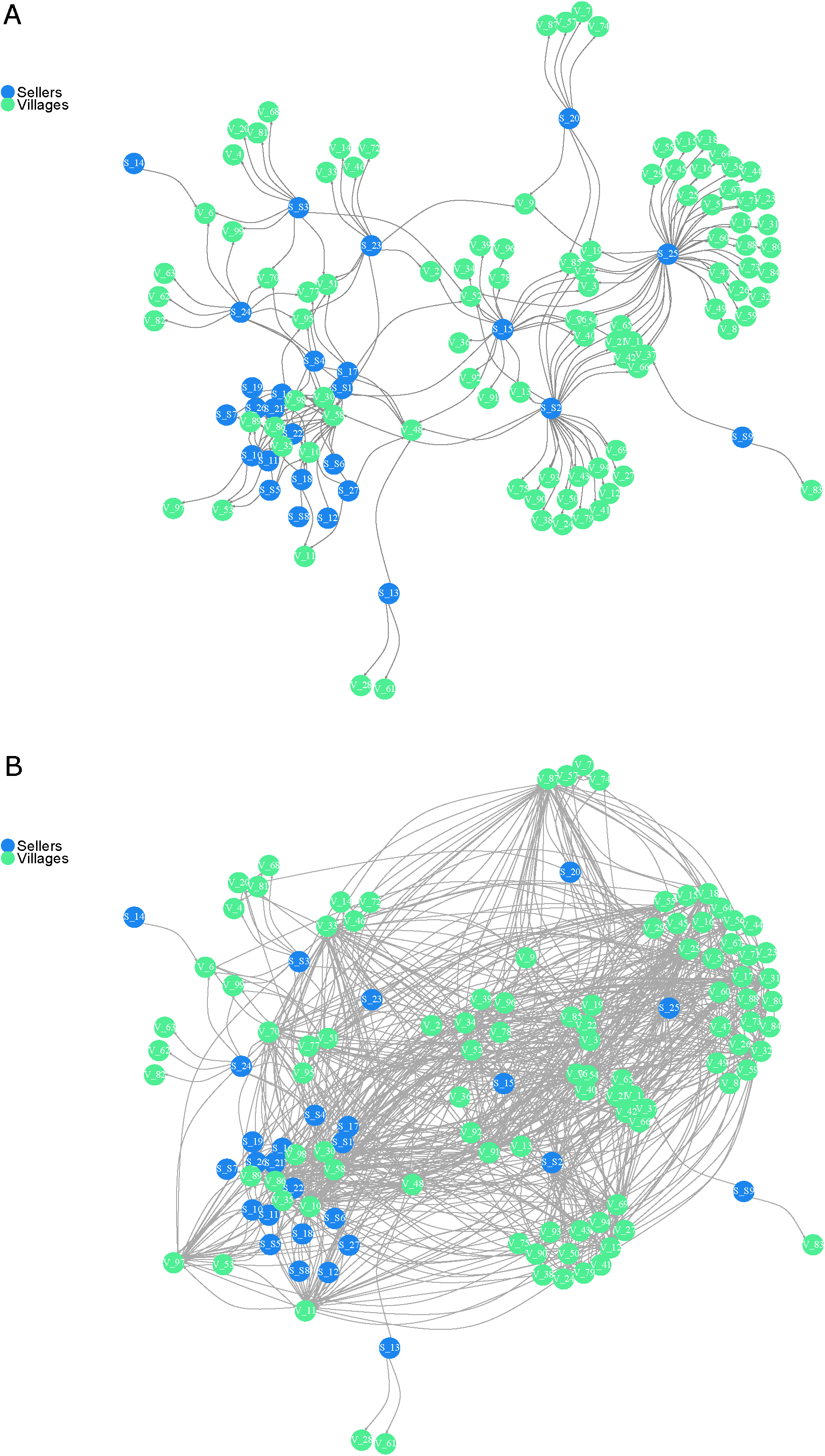
The network structure of sweetpotato vine transactions reported in the 2014 growing season in Northern Uganda with both sellers (darker nodes) and villages (lighter nodes). Links represent the occurrence of at least one transaction in the 2014 growing season. Note that the network layout is generated by the Fruchterman-Reingold force directed algorithm, which locates nodes with links closer together and those without links further apart, and not the geographic coordinates of villages. Plot A) represents empirically sampled seller-to-village links and, B) represents village-to-village links estimated as a function of inter-village distance.

**Figure 3.**
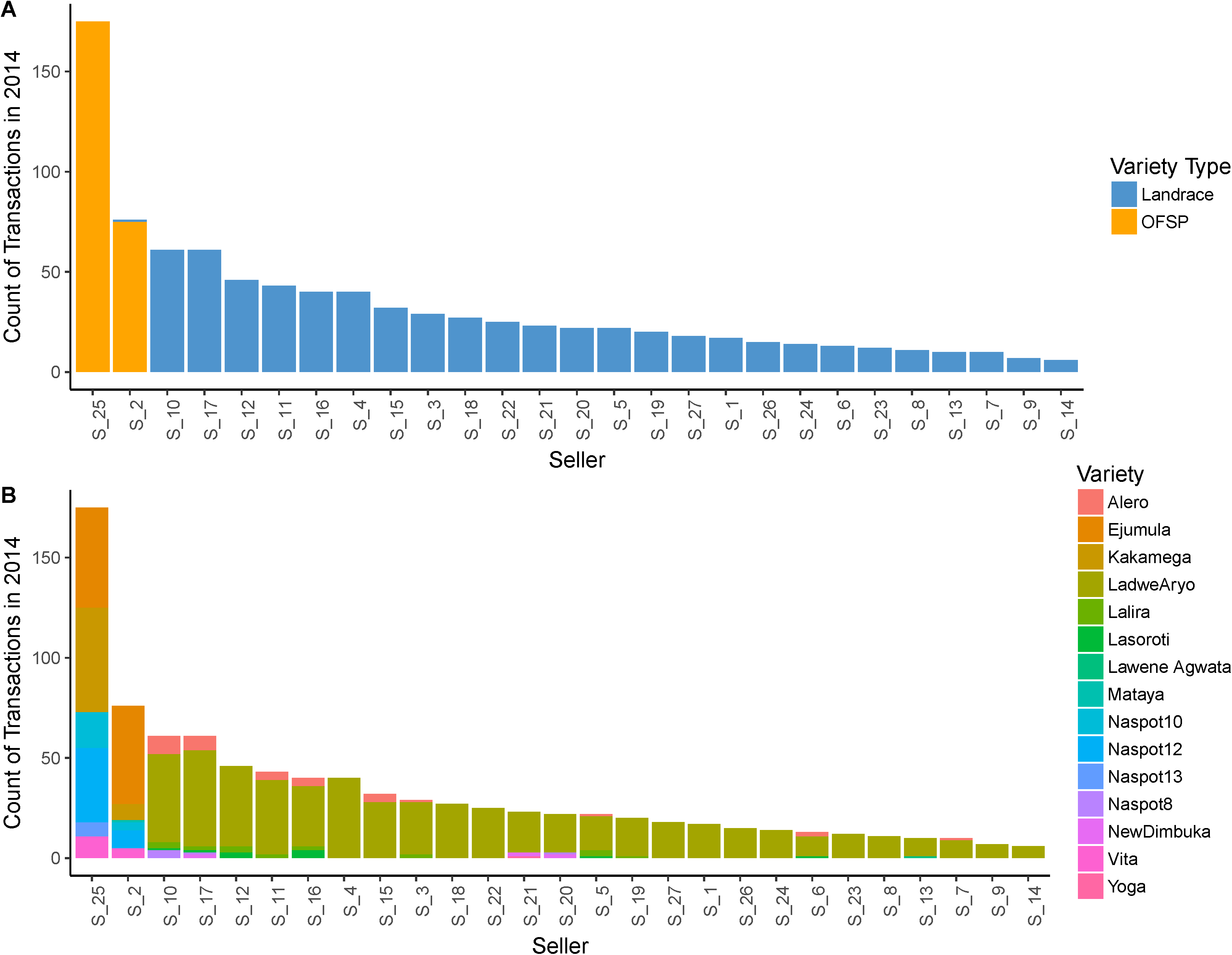
Bar chart indicating the reported transaction count per seller for A) varieties grouped as landrace or orange-fleshed sweetpotato (OFSP) varieties, and B) all individual varieties that were sold in 2014. Sellers were visited once per week during the growing season to obtain their reported transaction history for the previous week. Two sellers in the seed network sell OFSPs, with the rest selling only landraces. The landrace variety Ladwe Aryo is sold by 93% of sellers surveyed, with a higher number of transactions than any other variety in 2014.

### Variety Dissemination

A total of 15 cultivars were sold during the 2014 season (Fig. 3). These cultivars were a mix of landraces (all white-fleshed) and cultivars introduced by the national breeding program (Fig. 3). Six of these cultivars were OFSP cultivars, and were disseminated by only two sellers, in many individual transactions (Fig. 3). In comparison, the most common white-fleshed landrace, Ladwe Aryo, was sold by 25 distinct sellers in hundreds of transactions throughout the season. When the network is examined separately by variety, disaggregation becomes apparent (Fig. 4). Although both Ladwe Aryo and the OFSP cultivar, Ejumula, reached 51 villages each, only 8 of these villages were overlapping (Fig 4). Evaluating networks representing the distribution of the top eight varieties to villages in the network (Fig. 4) indicates that only a small number of sellers and villages were exchanging orange-fleshed varieties. It appears, from this survey, that most individuals from a single village only buy a single variety, even when they have access to multiple sellers.

**Figure 4.**
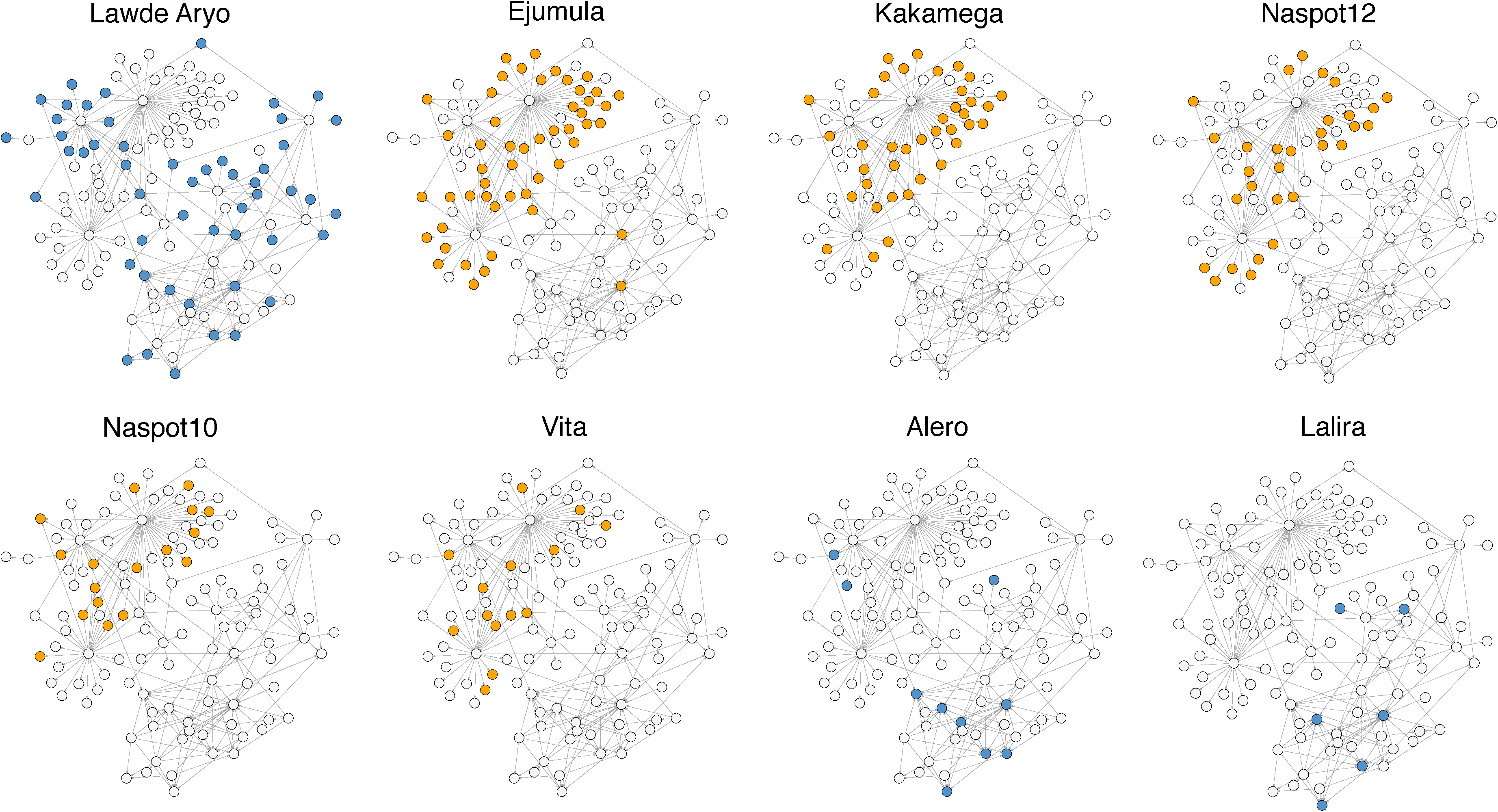
Networks of dissemination of the top eight cultivars sold during the 2014 growing season, in terms of number of transactions reported. All sellers and villages surveyed are represented in each network, while filled nodes represent sellers and villages involved in the sale or purchase of the specified cultivar. Unfilled villages did not access a given variety in 2014 through this surveyed seed network. Node shade indicates white-fleshed landraces (darker) and OFSP (lighter) cultivars.

### Experiment 1: The value of villages as risk-based surveillance locations

For the scenario where each node is an equally likely starting point for an epidemic, we evaluated the value of each village as a risk-based surveillance location. A village was considered a more effective location if only a small proportion of other nodes were colonized before the pathogen could be detected at that village. The village with the highest surveillance value was V_58, with the lowest mean number of villages affected by the time the pathogen would be detected there (Fig. 5). This village was close to the town seller, where many of the sellers sold their vines, likely accounting for high access to vines and a central position in the network.

**Figure 5.**
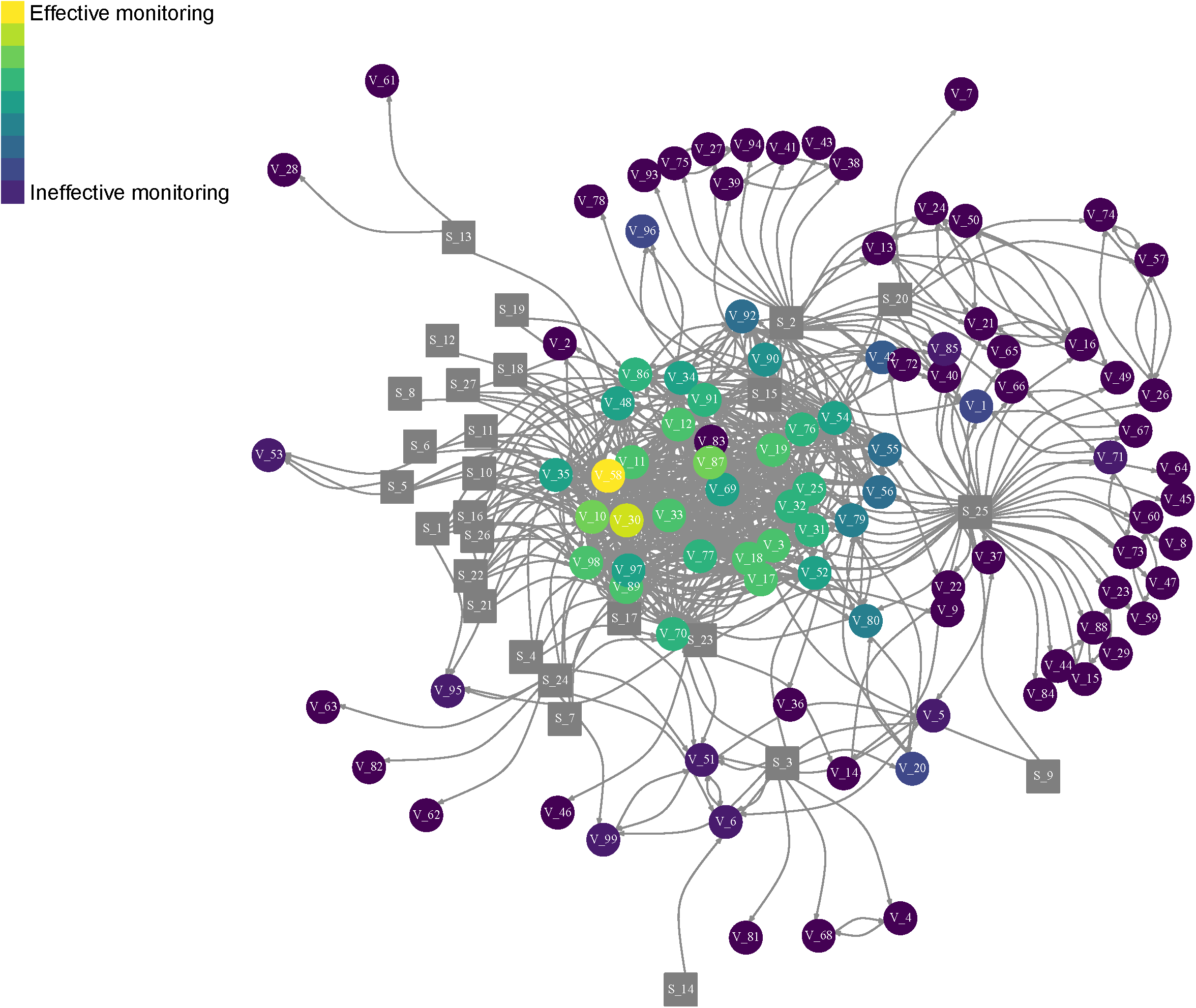
Network resulting from a simulation (500 realizations) where each node had equal likelihood of being the introduction point of an invasive pathogen. Node color indicates the total number of nodes that would be reached in the network before detection at that node. If few nodes were reached (lighter shading) the node was considered an effective monitoring location, while if many nodes were reached (darker shading) the node was considered ineffective for monitoring. Nodes labeled with an “S” and depicted with gray boxes are sellers, and circles (“V”) are villages. Only villages were considered in this analysis for their efficacy as sampling locations.

### Experiment 2: Modeling epidemic progress with variable starting sellers

The two sellers with the greatest out-degree for their product, S_25 (OFSP seller) and S_15 (landrace seller), were compared as starting nodes for an introduced epidemic simulated over 20 timesteps (Fig. 6). These sellers had an out-degree of 42 and 19, respectively. Infection starting with the landrace seller (S_15) approached epidemic saturation (where the maximum number of villages that can be reached, has been reached (50 of the 99 villages)). For the OFSP seller, saturation was at approximately 70 nodes out of 99. Overall, the mean number of nodes infected by time 20 was lower when infection started with the landrace seller (50.2 sd = 2.1) than for the OFSP seller (69.4 sd = 2.5). Disease progress, therefore, differed between starting nodes as well, with AUDPC being higher for infection starting with the OFSP seller (977, sd = 54.6) compared to starting with the landrace seller (697, sd = 57.2).

**Figure 6.**
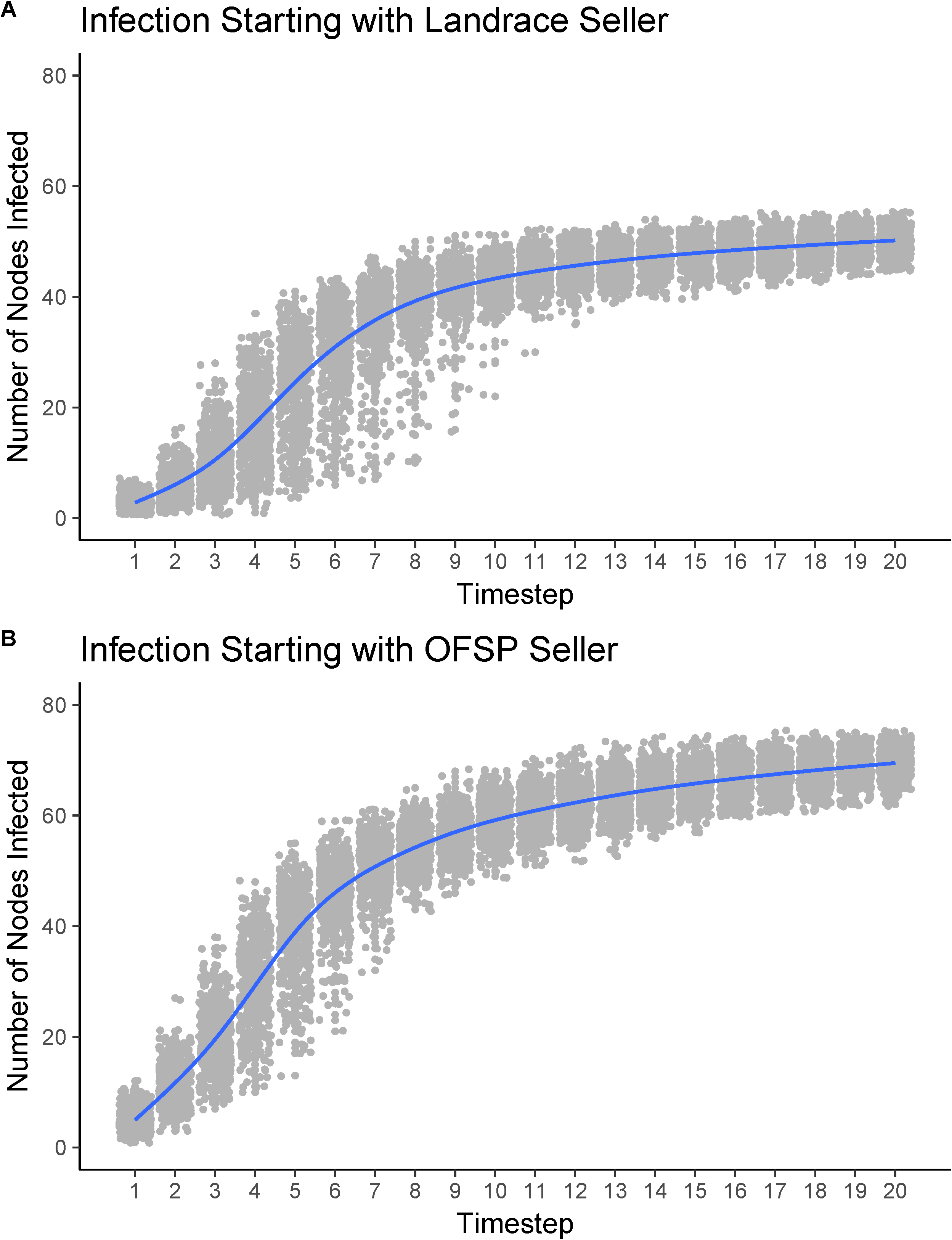
Epidemic progress through the seed network over 20 hypothetical time steps in 500 realizations (each realization indicated as a gray point) with the darker line indicating the smoothed conditional mean (local regression) of disease progress across realizations. Epidemics were modeled with two starting sellers, A) the seller with the highest out-degree of landrace sales (S_15), and b) the seller with the highest out-degree of orange-fleshed sweetpotato cultivar sales (S_25). Number of nodes infected represents the sum of the infection statuses of all nodes at each time step.

### Experiment 3: The influence of quarantine on infection dynamics

We evaluated the influence of quarantining villages so they cannot become infected or transmit infections to other villages, representing the common practical scenario of phytosanitary quarantine by regulatory agencies after the detection of pathogens in new regions. Nodes were selected based on node degree rank (with comparison to other methods of selecting nodes in the next step). Node degree for the 30 nodes selected as quarantine candidates ranged from 60-17 (with the highest selected first). Epidemic simulations, as previously described, were repeated for both sellers (S_25 and S_15), for each quarantine scenario based on degree (Fig. 7). For infection beginning with a landrace seller (Fig. 7a) the mean number of nodes infected by the end of the simulation (timestep 20) decreased by 6%, 16%, 28%, 46%, 66%, 71%, 78%, 79% and 81% for each quarantine treatment (quarantine of 10, 15, 20, 25, 30, 35, 40, 45, and 50 nodes, respectively) when compared to the no-quarantine control. The AUDPC decreased similarly with increasing number of quarantine nodes (Fig 7b).

**Figure 7.**
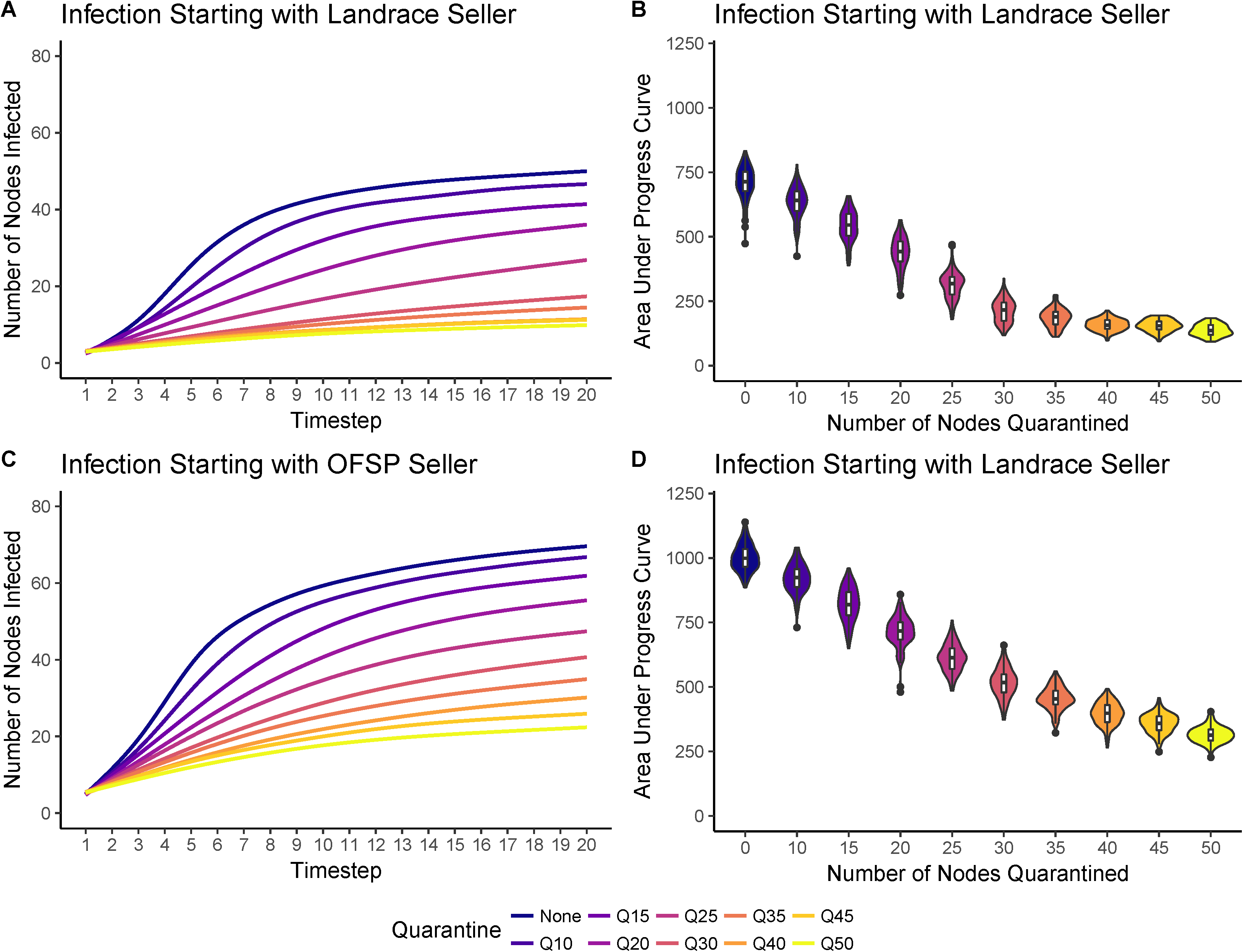
Disease progress over 20 time steps under specified quarantine regimes (None = no quarantine, and 10, 15, 20, 25, 30, 35, 40, 45, or 50 nodes quarantined). A) and C) Disease progress, for each quarantine status, with infection starting with the seller with the highest out-degree of landrace sales (S_15) in A, or the seller with highest out-degree of orange-fleshed sweetpotato sales (S_25) in C. Lines represent smoothed conditional means. B) and D) Area under the disease progress curve boxplot and distribution (violin plot) for each of the 500 realizations in A and C, respectively.

For the OFSP seller, the mean number of nodes infected at the end of time 20 decreased by 5%, 17%, 28%, 44%, 57%, 70%, 79%, 88%, and 94% for quarantine of 10, 15, 20, 25, 30, 35, 40, 45, and 50 nodes, respectively (Fig. 7c). Across treatments, final epidemic outcome was higher when infection starting with the OFSP seller (S_25), and although a similar effect of quarantine was observed, the percent control imposed by the treatment appeared to variable between sellers (S_17). To further explore whether the node degree of the starting seller was a main driver of the success of quarantine, quarantine efficacy, calculated as (AUDPC_no quarantine_ – AUDPC_quarantined_) / (AUDPC_no quarantine_), was examined for all sellers (Fig. S4) and accounted for ∼ 17% of the variation in AUDPC response.

We compared the selection of quarantine candidates based on node degree centrality (used in analysis described above) to closeness, betweenness, eigenvector centrality, the estimated risk-based surveillance score, and villages drawn at random, in terms of how they affected AUDPC outcomes across realizations (Figure 8). It was not surprising that, after quarantining at least 15 nodes, the “smart” selection criteria (centrality measures and monitoring efficiency) outperformed the “naive” selection criteria (randomly selecting villages to quarantine) (Supplemental Figs. 5 and 6). It was somewhat surprising, however, that this trend was not clear for the 10 node quarantine treatment, indicating that the invasion may be able to “overcome” quarantine of less than 10 nodes, even if the nodes are those with the highest centrality scores.

**Figure 8.**
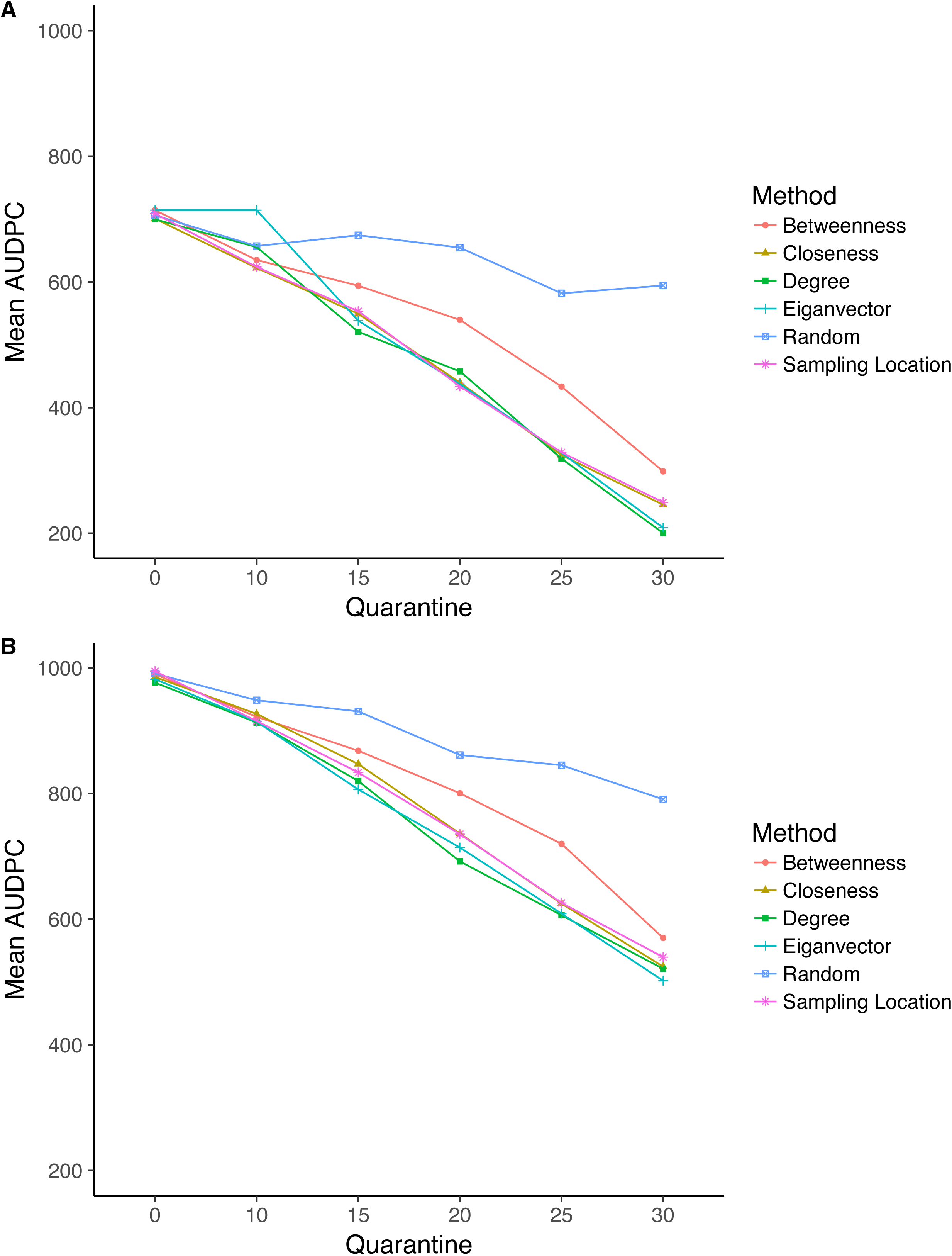
Mean AUDPC values for six quarantine scenarios (0 = no quarantine, and 10, 15, 20, 25, or 30 nodes quarantined, respectively) for infection starting with A) the seller with the highest out-degree of landrace sales (S_15), and B) the seller with the highest out-degree of orange-fleshed sweetpotato sales (S_25). Four different centrality measures were compared for their utility in selecting quarantine locations (betweenness, closeness, degree, and eigenvector). These were also compared to a scenario where nodes were selected at random for quarantine, and nodes that were selected as effective monitoring locations (“Sampling Location”) in the previously described simulation experiment (Fig. 5).

In this system, it appears that closeness, eigenvector, and degree centrality are equally good predictors of quarantine hubs and confer a similar reduction to AUDPC (Figure 8). This is consistent for each of the starting nodes. Interestingly, independent of starting node, betweenness centrality consistently resulted in a higher mean AUDPC when 20 and 25 nodes were quarantined.

### Sensitivity analysis and uncertainty quantification

Sensitivity analysis was performed to examine the influence of key parameters, describing the network and epidemic spread, on final epidemic outcomes. Both the exponent of the power law β and the threshold for link formation *t* influence the epidemic progress curve (Fig. 9). As β and *t* decrease, we see an increase in the number of nodes infected by the end of the epidemic, time 20 (Fig. 9). At time 20, with β = 0.5 and *t =* 0.001 all villages are infected, in each of the 500 realizations. Alternatively, when β = 2.5 and *t =* 0.02, little or no epidemic progress is made. The same trend is apparent for infection starting with each of the two sellers (Supplemental Fig. S7). These dramatic differences are likely due to the number of links formed between villages with each parameter set. With the network highly saturated with links, as is the case with β = 0.5 and *t =* 0.001, the epidemic can move across the system rapidly. A similar trend was also seen when the probability of transmission (*P*_*t*_) varied from 0.1 to 0.05 and 0.15 (Supplemental Fig. S8).

**Figure 9.**
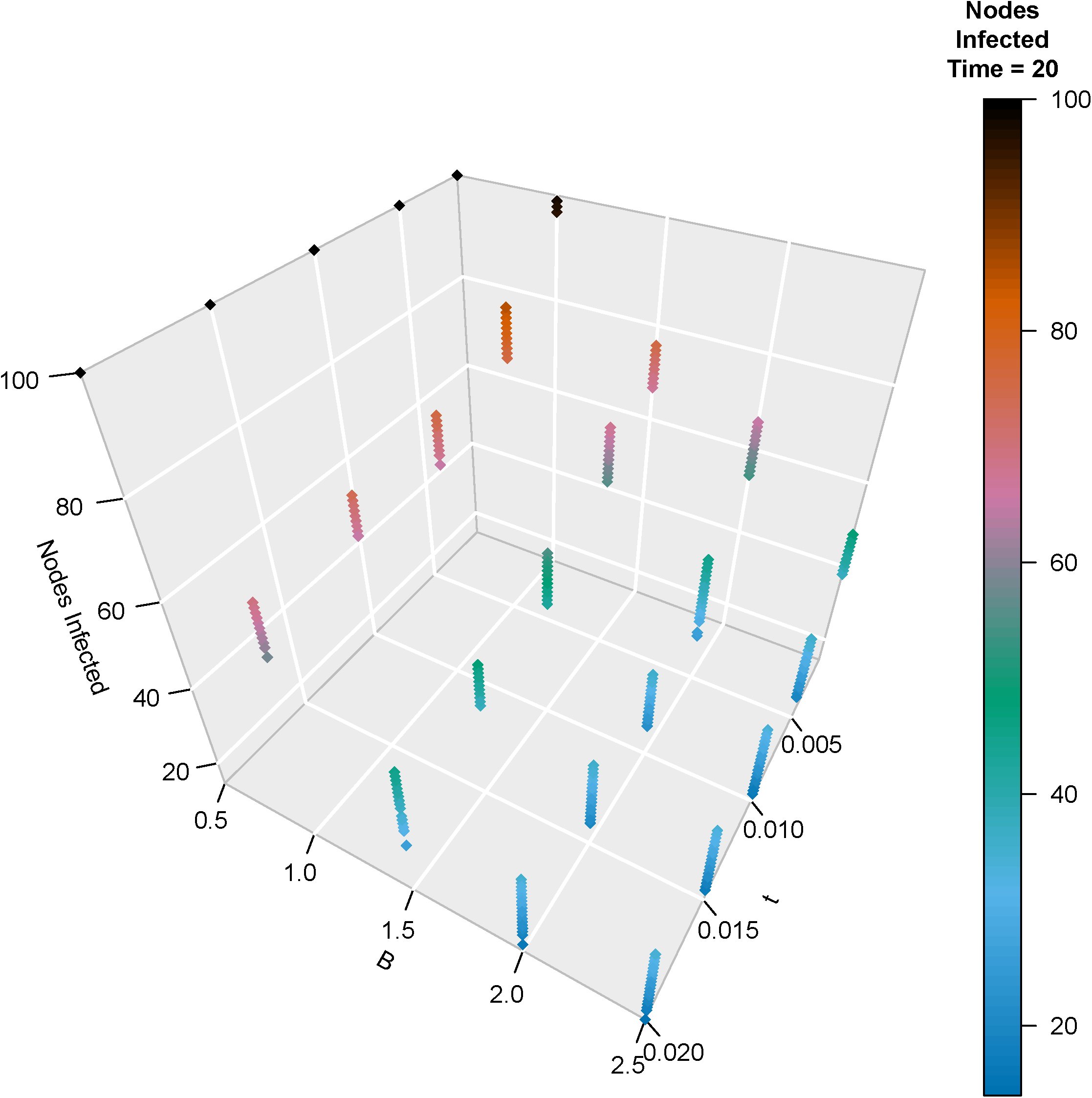
Uncertainty quantification for the number of nodes infected at time step 20 across 500 realizations for a range of values for parameters β (0.5-2.5) and *t* (.001-.02). β is the exponent of the power law equation, and *t* is a threshold applied to the power-law transformed distance village distance matrix. β and *t* were used to evaluate seed transaction links between villages, and therefore influenced disease spread. The simulated epidemic started at node 15 (S_15).Similar analyses for other starting nodes and a range of transmission rates (*P*_*t*_) were also performed (Figs. S5 and S6).

## Discussion

In this study, we layered empirical survey data with geo-spatial data to construct a network model of the sweetpotato seed system in Northern Uganda. Using this model, we simulated epidemics under several scenarios and identified key villages (nodes) in the region for both monitoring and for the deployment of management tactics, like quarantine. Epidemic starting point influenced not only epidemic outcome, but also the efficacy of quarantine interventions for limiting disease spread. In this system we compared several centrality measures for their utility to limit epidemic spread when placed under quarantine. We found that node degree performed as well as or better than centrality measures that take into account the broader network topology (eigenvector centrality, betweenness centrality, and closeness centrality), a finding with practical management implications.

The analysis of network topologies presented here shows how sales of OFSP varieties and landrace varieties differed. When vine distribution was disaggregated by variety, it is clear that most cultivars are not well disseminated by seller-to-village links (Fig 4). We found that a single white-fleshed landrace variety, Ladwe Aryo, dominated landrace sales in this season, while a single OFSP vine, Ejumula, dominated OFSP sales (Figs. 3 and 4). Interestingly, there is little overlap between the villages where farmers bought these varieties. Based on the observed data, we cannot determine whether preference or availability of planting material of OFSP varieties drove sales, or a combination of the two. This distinction is important, however, because much effort has been made to promote improved OFSP varieties in recent decades (Low et al 2017). In a study in 2015 (Obong et al. 2017), 89% of 51 multiplier fields sampled in the Gulu region during the 2015 off-season grew local landraces, with the remainder of area planted to improved varieties. This proportionally low rate of adoption might be attributed to the very recent introduction of OFSPs into the region (with promotion by HarvestPlus and affiliated NGOs), and may increase in coming years if these varieties become accepted by farmers and there remains an adequate local supply of vines provided by these informal sellers. Seed exchange is often highly related to kinship ties, language, and social organization patterns (Abizaid et al. 2016; Labeyrie et al. 2016; Perales et al. 2005; Westengen et al. 2014) and future research to better model the influence of social structure on variety diffusion in this seed system network would be useful.

Our findings are consistent with previous studies indicating that the epidemic starting point influences both epidemic progress and epidemic outcome (Pautasso 2015; Pautasso et al. 2010). This was illustrated in experiments 2 and 3 where we found that when the epidemic started with the highest out-degree OFSP seller, when compared to the highest out-degree landrace seller, more villages in the landscape became infected by the conclusion of the epidemic. It appears that one main driver of this phenomenon is the out-degree of the “starting seller” (or the seller with infected seed material). We also found that the epidemic starting seller influenced the efficacy of quarantine interventions, with a lower percent control conferred by the same quarantine treatment when infection started with the OFSP seller, compared to a landrace seller, up to a certain number of nodes quarantined (30) after which the trend was reversed. This indicates that not only does the introduction point of a pathogen in this network impact the risk of spread through the network, but it also influences the efficacy of management strategies.Similar methods have also been used to simulate variety diffusion through seed systems (Pautasso 2015). Future studies should assess the rate of adoption over time in the landscape, to determine the implications of the models of spread that we presented for each type of variety (Fig 5).

In experiment 1 we characterized villages in Northern Uganda for their utility as pathogen surveillance locations, based on a set of simulations where all nodes had an equal chance of being the point of entry into the network. Villages were identified as potential risk-based surveillance targets based on the observed frequency, in simulations, that the pathogen would be detected in these locations before substantial parts of the rest of the network become colonized. This method can be used to identify “high utility” sentinel locations to focus sampling efforts for new and emerging pathogens, a critical effort to avoid large-scale epidemics. This method of prescribing surveillance locations can complement new field-based diagnostic technologies, such as loop-mediated amplification (LAMP) assays (Yasuhara-Bell et al. 2016) or smartphone-assisted crop disease image detection (Mohanty et al. 2016), which are becoming increasingly available to practitioners and have the potential for rapid on-site detection of viruses and other pathogens. The method used here, applied to a region in northern Uganda, can support pathogen surveillance efforts on a national or greater scale.

Centrality measures are important indicators of risk within epidemic networks (Banks et al. 2015; Holme 2017; Kiss et al. 2006). There is a trade-off of high centrality within this seed network. That is, it is favorable to be a village with high degree (many links) because of the increased availability of a diversity of cultivars (like OFSP varieties), from more sellers.Availability is often defined as the physical presence of seed at planting time within a reasonable proximity of farmers (McGuire and Sperling 2011), and is a component of seed security for a village. On the other hand, high connectivity makes a village more susceptible to pathogen invasion and increases its likelihood of serving as an epidemic “super-spreader”, as illustrated by our quarantine experiments (Fig 7) where, after high-degree villages were quarantined, the epidemic was successfully reduced. Node degree was a good measure for the informed selection of quarantine locations (Fig. 8). The utility of node degree for prioritization is consistent with literature on pathogen spread in animal trade networks, where centrality measures (such as degree) have been significantly associated with on-farm disease levels and disease progress (Kiss et al. 2006; Lee et al. 2017; Salines et al. 2018).

We compared the use of node degree, with node betweenness, closeness, and eigenvector centrality, and found each performing similarly, with betweenness conferring the least control.Practically, this is an important finding for the mitigation of invasive pathogens in plant exchange networks, as node degree is one of the simplest centrality measures to collect (Christley et al. 2005) and confirms findings from human epidemiology literature (Lloyd-Smith et al. 2005). In times of epidemic emergence, these high degree villages should be the first to be targeted with control strategies, such as quarantine in the form of phytosanitary regulation.Similar methods to those described here may be utilized to target villages for development projects that aim to disseminate varieties to key hubs and maximize their distribution. However, betweenness may prove to be more important for identifying key nodes in some other types of seed systems, in which there are many nearly separate modules with a small number of links between them. In such seed systems, the nodes with high betweenness centrality that bridge these modules might be particularly important.

Although quarantine was effective at *slowing* epidemic progress when sufficient nodes were included, here we found that quarantine was not sufficient (at < 50 nodes quarantined) for *halting* epidemic spread under any scenario. This property of rapid disease spread is consistent with other scale-free networks, where high-degree “super-spreaders” can rapidly transmit disease to other nodes in the network in a small number of steps (Banks et al. 2015; Jeger et al. 2007; Lloyd-Smith et al. 2005).

Future research should consider incorporating the influence of the deployment of other components of an integrated seed health strategy (Thomas-Sharma et al. 2017) into the model for epidemic mitigation, such as positive selection, resistant varieties, clean seed, and education about disease progress and management strategies. Integrated seed health may be easier and more cost effective to deploy than strict phytosanitary restrictions or introducing complete ‘quarantine’ of villages in these systems, where ethnobiological associations are major drivers of exchange between villages, and access to certified clean seed may be minimal to non-existent.

It is important to note that this study was based on data limited to sellers that participated in weekly monitoring in the Gulu region of Northern Uganda. Although this dataset was expansive, there may have been other sellers or sources of vines that were not captured. In addition, there are other villages with sweetpotato fields in the region of study that were not included in this analysis because they were not associated with a buyer in the dataset. These fields could also be a source of disease. Furthermore, links in this study were unweighted and fixed through time. It may be valuable to examine the influence of a weighted, dynamic contact structure on epidemic progress in such seed networks. To our knowledge, this has not been examined in the literature. The model used here is a susceptible-infected (SI) model, where once nodes become infected, they remain infected (and infectious) through the duration of the time course. In human epidemiology, diseases that follow an SI model are known to be difficult to control. Because there may be options for epidemic recovery in this sweetpotato seed system (through reversion, roguing, or positive selection), future studies may include a recovery term in the model, which would result in more effective quarantine measures. Another outstanding question is the influence of multiple starting points on epidemic progress and outcome.

Understanding the dynamics of epidemics in seed systems is critical for effective pathogen monitoring, risk assessment, and epidemic management (Buddenhagen et al. 2017; Harwood et al. 2009; Shaw and Pautasso 2014). Although some plant disease literature explores this topic, research on real world seed network epidemics remains limited. Results from these studies can be strategically used to prioritize surveillance efforts and to disseminate new varieties in informal seed systems. Future surveys that include questions about social ties and the movement of information among farmers would support better models of variety adoption and distribution in this system. Next research steps will include more finely parameterizing transmission patterns, including the impact of variety resistance and vector biology, and modeling system adaptation to sustained exogenous shocks and stressors. There is the potential to include data about known yield degeneration rates and known environmental conditions to predict regional yield loss in the case of pathogen introduction. Understanding these system components supports better strategies for seed system development.

## Acknowledgements

This work received institutional review board approval through the University of Florida IRB-02 #IRB201700024. We confirm farmer participation was voluntary and that personal and demographic information was protected. This research was undertaken as part of, and funded by, the CGIAR Research Program on Roots, Tubers and Bananas (RTB) and supported by CGIAR Fund Donors http://www.cgiar.org/about-us/governing-2010-june-2016/cgiar-fund/fund-donors-2/, The Bill and Melinda Gates Foundation (OPP1080975), US NSF Grant EF-0525712 as part of the joint NSF-NIH Ecology of Infectious Disease program, US NSF Grant DEB-0516046, and the University of Florida. The authors confirm they have no conflicts of interest.

## Literature Cited

Abay, F., de Boef, W., and Bjørnstad, Å. 2011. Network analysis of barley seed flows in Tigray, Ethiopia: supporting the design of strategies that contribute to on-farm management of plant genetic resources. Plant Genetic Resources 9:495–505.

Abizaid, C., Coomes, O. T., and Perrault-Archambault, M. 2016. Seed sharing in Amazonian indigenous rain forest communities: a social network analysis in three Achuar villages, Peru. Human Ecology 44:577–594.

Adikini, S., Mukasa, S. B., Mwanga, R. O. M., and Gibson, R. W. 2015. Sweet potato cultivar degeneration rate under high and low sweet potato virus disease pressure zones in Uganda. Canadian Journal of Plant Pathology 37:136–147.

Almekinders, C. J. M., Louwaars, N. P., and Debruijn, G. H. 1994. Local seed systems and their importance for an improved seed supply in developing countries. Euphytica 78:207–216.

Banks, N. C., Paini, D. R., Bayliss, K. L., and Hodda, M. 2015. The role of global trade and transport network topology in the human-mediated dispersal of alien species. Ecol Lett 18:188–199.

Barabasi, A.-L., and Albert, R. 1999. Emergence of scaling in random networks. Science 286:509–512.

Behrman, J. 2011. HarvestPlus Reaching End Users (REU) Orange-Fleshed Sweet Potato (OFSP) Project: Report of Qualitive Findings from Uganda. International Food Policy Research Institute 1–29.

Buddenhagen, C. E., Hernandez Nopsa, J. F., Andersen, K. F., Andrade-Piedra, J., Forbes, G. A., Kromann, P., Thomas-Sharma, S., Useche, P., and Garrett, K. A. 2017. Epidemic Network Analysis for Mitigation of Invasive Pathogens in Seed Systems: Potato in Ecuador. Phytopathology 107:1209–1218.

Christley, R. M., Pinchbeck, G. L., Bowers, R. G., Clancy, D., French, N. P., Bennett, R., and Turner, J. 2005. Infection in social networks: using network analysis to identify high-risk individuals. Am J Epidemiol 162:1024–1031.

Csárdi, G., and Nepusz, T. 2006. The igraph software package for complex network research. InterJournal, Complex Systems 1695:1–9.

De Groote, H., Oloo, F., Tongruksawattana, S., and Das, B. 2016. Community-survey based assessment of the geographic distribution and impact of maize lethal necrosis (MLN) disease in Kenya. Crop Protection 82:30–35.

Garrett, K. A. 2012. Information networks for disease: commonalities in human management networks and within-host signalling networks. European Journal of Plant Pathology 133:75–88.

Garrett, K. A., Andersen, K. F., Asche, F., Bowden, R. L., Forbes, G. A., Kulakow, P. A., and Zhou, B. 2017. Resistance genes in global crop breeding networks. Phytopathology 107:1268–1278.

Gibson, R. 2013. How sweet potato varieties are distributed in Uganda: actors, constraints and opportunities. Food Security 5:781–791.

Gibson, R. W., and Kreuze, J. F. 2015. Degeneration in sweetpotato due to viruses, virus-cleaned planting material and reversion: a review. Plant Pathology 64:1–15.

Gibson, R. W., Namanda, S., and Sindi, K. 2011. Sweetpotato seed systems in East Africa. African Crop Science Conference Proceedings 10:449–451.

Gildemacher, P. R., Demo, P., Barker, I., Kaguongo, W., Woldegiorgis, G., Wagoire, W. W., Wakahiu, M., Leeuwis, C., and Struik, P. C. 2009. A description of seed potato systems in Kenya, Uganda and Ethiopia. American Journal of Potato Research 86:373–382.

Graziosi, I., Minato, N., Alvarez, E., Ngo, D. T., Hoat, T. X., Aye, T. M., Pardo, J. M., Wongtiem, P., and Wyckhuys, K. A. 2016. Emerging pests and diseases of South-east Asian cassava: a comprehensive evaluation of geographic priorities, management options and research needs. Pest Management Science 72:1071–1089.

Harwood, T. D., Xu, X., Pautasso, M., Jeger, M. J., and Shaw, M. W. 2009. Epidemiological risk assessment using linked network and grid based modelling: Phytophthora ramorum and Phytophthora kernoviae in the UK. Ecological Modeling 220:3353–3361.

Hernandez Nopsa, J. F., Daglish, G. J., Hagstrum, D. W., Leslie, J. F., Phillips, T. W., Scoglio, C., Thomas-Sharma, S., Walter, G. H., and Garrett, K. A. 2015. Ecological networks in stored grain: Key postharvest nodes for emerging pests, pathogens, and mycotoxins. BioScience 65:985–1002.

Hilker, F. M., Allen, L. J. S., Bokil, V. A., Briggs, C. J., Feng, Z., Garrett, K. A., Gross, L. J., Hamelin, F. M., Jeger, M. J., Manore, C. A., Power, A. G., Redinbaugh, M. G., Rua, M. A., and Cunniffe, N. J. 2017. Modeling virus coinfection to inform management of maize lethal necrosis in Kenya. Phytopathology 107:1095–1108.

Holme, P. 2017. Three faces of node importance in network epidemiology: Exact results for small graphs. Phys Rev E 96:062305.

Jeger, M. J., Pautasso, M., Holdenrieder, O., and Shaw, M. W. 2007. Modelling disease spread and control in networks: implications for plant sciences. New Phytol 174:279–297.

Johnson, A. C., and Gurr, G. M. 2016. Invertebrate pests and diseases of sweetpotato (Ipomoea batatas): a review and identification of research priorities for smallholder production. Annals of Applied Biology 168:291–320.

Karyeija, R. F., Gibson, R. W., and Valkonen, J. P. T. 1998. The significance of sweet potato feathery mottle virus in subsistence sweet potato production in Africa. Plant Disease 82:4–15.

Kiss, I. Z., Green, D. M., and Kao, R. R. 2006. The network of sheep movements within Great Britain: Network properties and their implications for infectious disease spread. J R Soc Interface 3:669–677.

Labeyrie, V., Thomas, M., Muthamia, Z. K., and Leclerc, C. 2016. Seed exchange networks, ethnicity, and sorghum diversity. Proc Natl Acad Sci U S A 113:98–103.

Lee, K., Polson, D., Lowe, E., Main, R., Holtkamp, D., and Martinez-Lopez, B. 2017. Unraveling the contact patterns and network structure of pig shipments in the United States and its association with porcine reproductive and respiratory syndrome virus (PRRSV) outbreaks. Prev Vet Med 138:113–123.

Lloyd-Smith, J. O., Schreiber, S. J., Kopp, P. E., and Getz, W. M. 2005. Superspreading and the effect of individual variation on disease emergence. Nature 438:355–359.

Mahuku, G., Lockhart, B. E., Wanjala, B., Jones, M. W., Kimunye, J. N., Stewart, L. R., Cassone, B. J., Sevgan, S., Nyasani, J. O., Kusia, E., Kumar, P. L., Niblett, C. L., Kiggundu, A., Asea, G., Pappu, H. R., Wangai, A., Prasanna, B. M., and Redinbaugh, M. G. 2015. Maize Lethal Necrosis (MLN), an emerging threat to maize-based food security in Sub-Saharan Africa. Phytopathology 105:956–965.

McGuire, S., and Sperling, L. 2011. The links between food security and seed security: facts and fiction that guide response. Development in Practice 21:493–508.

McGuire, S., and Sperling, L. 2013. Making seed systems more resilient to stress. Global Environmental Change 23:644–653.

McQuaid, C. F., van den Bosch, F., Szyniszewska, A., Alicai, T., Pariyo, A., Chikoti, P. C., and Gilligan, C. A. 2017. Spatial dynamics and control of a crop pathogen with mixed-mode transmission. PLoS Comput Biol 13:e1005654.

Mohanty, S. P., Hughes, D. P., and Salathe, M. 2016. Using deep learning for image-based plant disease detection. Front Plant Sci 7:1419.

Moslonka-Lefebvre, M., Ann Finley, I. D., Dehnen-Schmutz, K., Harwood, T., Jeger, M. J., Xu, X., Holdenrieder, O., and Pautasso, M. 2011. Networks in plant epidemiology: from genes to landscapes, countries, and continents. Phytopathology 101:392–403.

Nelson, M. F., and Bone, C. E. 2015. Effectiveness of dynamic quarantines against pathogen spread in models of the horticultural trade network. Ecological Complexity 24:14–28.

Obong, Y., Omony, T., Rachkara, P., and Gibson, R. W. 2017. Disseminating modern sweetpotato varieties using participatory variety demonstration trials on informal nodal multipliers’ fields at hub locations. Journal of Crop Improvement:1–18.

Pautasso, M. 2015. Network simulations to study seed exchange for agrobiodiversity conservation. Agronomy for Sustainable Development 35:145–150.

Pautasso, M., and Jeger, M. J. 2008. Epidemic threshold and network structure: The interplay of probability of transmission and of persistence in small-size directed networks. Ecological Complexity 5:1–8.

Pautasso, M., Moslonka-Lefebvre, M., and Jeger, M. J. 2010. The number of links to and from the starting node as a predictor of epidemic size in small-size directed networks. Ecological Complexity 7:424–432.

Pautasso, M., Aistara, G., Barnaud, A., Caillon, S., Clouvel, P., Coomes, O. T., Delêtre, M., Demeulenaere, E., De Santis, P., Döring, T., Eloy, L., Emperaire, L., Garine, E., Goldringer, I., Jarvis, D., Joly, H. I., Leclerc, C., Louafi, S., Martin, P., Massol, F., McGuire, S., McKey, D., Padoch, C., Soler, C., Thomas, M., and Tramontini, S. 2013. Seed exchange networks for agrobiodiversity conservation. A review. Agron. Sustain. Dev. 33:151–175.

Perales, H. R., Benz, B. F., and Stephen, B. B. 2005. Maize diversity and ethnolinguistic diversity in Chiapas, Mexico. Proc Natl Acad Sci 102:949–953.

Pusadee, T., Jamjod, S., Chiang, Y.-C., and Rerkasem, B. 2009. Genetic structure and isolation by distance in a landrace of Thai rice. Proc Natl Acad Sci 106:13880–13885.

Rachkara, P., Phillips, D. P., Kalule, S. W., and Gibson, R. W. 2017. Innovative and beneficial informal sweetpotato seed private enterprise in northern Uganda. Food Security 9:595–610.

Salines, M., Andraud, M., and Rose, N. 2018. Combining network analysis with epidemiological data to inform risk-based surveillance: Application to hepatitis E virus (HEV) in pigs. Prev Vet Med 149:125–131.

Sanatkar, M. R., Scoglio, C., Natarajan, B., Isard, S. A., and Garrett, K. A. 2015. History, epidemic evolution, and model burn-in for a network of annual invasion: Soybean rust. Phytopathology 105:947–955.

Shaw, M. W., and Pautasso, M. 2014. Networks and plant disease management: concepts and applications. Annu Rev Phytopathol 52:477–493.

Silk, M. J., Croft, D. P., Delahay, R. J., Hodgson, D. J., Boots, M., Weber, N., and McDonald, R.A. 2017. Using social network measures in wildlife disease ecology, epidemiology, and management. Bioscience 67:245–257.

Sperling, L. 2008. When disaster strikes: a guide to assessing seed system security.

Sutrave, S., Scoglio, C., Isard, S. A., Hutchinson, J. M. S., and Garrett, K. A. 2012. Identifying highly connected counties compensates for resource limitations when evaluating national spread of an invasive pathogen. PLoS ONE 7:e37793.

Thomas-Sharma, S., Andrade-Piedra, J., Carvajal Yepes, M., Hernandez Nopsa, J., Jeger, M., Jones, R., Kromann, P., Legg, J., Yuen, J., Forbes, G., and Garrett, K. A. 2017. A risk assessment framework for seed degeneration: Informing an integrated seed health strategy for vegetatively-propagated crops. Phytopathology 107:1123–1135.

Thomas-Sharma, S., Abdurahman, A., Alic, S., Andrade-Piedrad, J. L., Bao, S., Charkowski, A. O., Crook, D., Kadian, M., Kromann, P., Struik, P. C., Torrance, L., Garrett, K. A., and Forbes, G. A. 2016. Seed degeneration in potato: the need for an integrated seed health strategy to mitigate the problem in developing countries. Plant Pathology 65:3–16.

Wang, H., Cui, X., Wang, X., Liu, S., and Zhou, X. 2016. First Report of Sri Lankan cassava mosaic virus Infecting Cassava in Cambodia. Plant Disease 100:1029.

Wangai, A. W., Redinbaugh, M. G., Kinyua, M., Miano, D. W., Leley, P. K., Kasina, M., Mahuku, G., Scheets, K., and Jeffers, D. 2012. First Report of Maize chlorotic mottle virus and Maize Lethal Necrosis in Kenya. Plant Disease 96:1582.

Westengen, O. T., Okongo, M. A., Onek, L., Berg, T., Upadhyaya, H., Birkeland, S., Kaur Khalsa, S. D., Ring, K. H., Stenseth, N. C., and Brysting, A. K. 2014. Ethnolinguistic structuring of sorghum genetic diversity in Africa and the role of local seed systems. Proc Natl Acad Sci U S A 111:14100–14105.

Yasuhara-Bell, J., de Silva, A., Heuchelin, S. A., Chaky, J. L., and Alvarez, A. M. 2016. Detection of Goss’s Wilt Pathogen Clavibacter michiganensis subsp. nebraskensis in Maize by Loop-Mediated Amplification. Phytopathology 106:226–235.

